# Synthetic whole-slide image tile generation with gene expression profiles infused deep generative models

**DOI:** 10.1101/2022.12.16.520705

**Authors:** Francisco Carrillo-Perez, Marija Pizurica, Michael G. Ozawa, Hannes Vogel, Robert B. West, Christina S. Kong, Luis Javier Herrera, Jeanne Shen, Olivier Gevaert

**Affiliations:** Stanford Center for Biomedical Informatics Research (BMIR), Stanford University, School of Medicine, 1265 Welch Rd, Stanford, 94305-547, CA, USA; Department of Architecture and Computer Technology (ATC), University of Granada, C. Periodista Daniel Saucedo Aranda, s/n, Granada, 18014, Granada, Spain; Internet technology and Data science Lab (IDLab), Ghent University, Technologiepark-Zwijnaarde 126, Gent, 9052, Gent, Belgium; Department of Pathology, Stanford University School of Medicine, 300 Pasteur Dr, Palo Alto, 94304, CA, USA; Department of Biomedical Data Science, Stanford University, School of Medicine, Medical School Office Building (MSOB), 1265 Welch Rd, Stanford, 94305-547, CA, USA

## Abstract

The acquisition of multi-modal biological data for the same sample, such as RNA sequencing and whole slide imaging (WSI), has increased in recent years, enabling studying human biology from multiple angles. However, despite these emerging multi-modal efforts, for the majority of studies only one modality is typically available, mostly due to financial or logistical constraints. Given these difficulties, multi-modal data imputation and multi-modal synthetic data generation are appealing as a solution for the multi-modal data scarcity problem. Currently, most studies focus on generating a single modality (e.g. WSI), without leveraging the information provided by additional data modalities (e.g. gene expression profiles). In this work, we propose an approach to generate WSI tiles by using deep generative models infused with matched gene expression profiles. First, we train a variational autoencoder (VAE) that learns a latent, lower dimensional representation of multi-tissue gene expression profiles. Then, we use this representation to infuse generative adversarial networks (GAN) that generate lung and brain cortex tissue tiles, resulting in a new model that we call RNA-GAN. Tiles generated by RNA-GAN were preferred by expert pathologists in comparison to tiles generated using traditional GANs and in addition, RNA-GAN needs fewer training epochs to generate high-quality tiles. Finally, RNA-GAN was able to generalize to gene expression profiles outside of the training set, showing imputation capabilities. A web-based quiz is available for users to play a game distinguishing real and synthetic tiles: https://rna-gan.stanford.edu/ and the code for RNA-GAN is available here: https://github.com/gevaertlab/RNA-GAN.

## Introduction

Biomedical data has become increasingly multi-modal, which has allowed us to better capture the complexity of biological processes. In the multi-modal setting, several technologies are used to obtain data from the same patient, providing a richer representation of their biological status and disease state. In current clinical practice, often demographic, clinical, molecular and imaging data are collected on patients. Making these data modalities available helps advancing the goals of precision medicine [1, 2]. For example, DNA and RNA-sequencing are now widely used for the characterization of cancer patients [3, 4]. Somatic mutation and gene expression profiles can be used to improve diagnosis, define disease subtypes, and determine the treatment regimen for cancer patients [5, 6]. Similarly, in pathology, tissue slides are the cornerstone for a variety of tasks. This includes primary diagnosis based on visual examinations by pathologists as well as treatment recommendations based on insights revealed by e.g. immunohistochemistry stains. [7]. Specifically for oncology, tissue slides are a valuable resource to observe morphological and texture changes, which reflect the tumor and its microenvironment [7–9]. Since the digitization of tissue slides to whole slide image (WSI) data, they have become a key data source for training deep learning models in a wide range of clinically relevant endpoints [10].

In particular, the relationship between genomic features and WSI image features has recently been demonstrated, with several studies showing that these two modalities are complementary. For example, morphological features from WSI data have been shown to associate with genomic mutations, gene expression profiles, and methylation patterns [5, 11, 12]. Moreover, studies have shown that the integration of both modalities leads to an improvement in the performance of machine learning models for diagnostic and prognostic tasks in cancer [5, 13–16].

However, both modalities are not always available due to financial or logistical constraints. For example, the Genome Express Omnibus (GEO) database [17] has numerous RNA-Seq datasets available, but few datasets have the corresponding WSI images. Similarly, most medical centers have large archives of tissue slides, but not yet the means to generate matched gene expression data. New multi-modal datasets are being created to deal with these issues [18], yet the problem still occurs for most clinical data sets. Thus, opportunities for training models that require multi-modal data are missed, slowing down progress in advancing precision medicine [19, 20].

Data scarcity is a concerning problem in the machine learning community, especially in the context of recent successes for non-medical applications where huge amounts of data are available [21, 22]. Specifically in biomedical problems, large and diverse cohorts are necessary to develop accurate clinical decision support systems that depend on machine learning algorithms [23]. To overcome the scarcity of heterogeneous annotated data in real-world biomedical settings, synthetic data is increasingly being considered [24]. Generative models, which impute synthetic data that are indistinguishable from real data, potentially offer a solution to deal with this issue. Within generative models, Generative Adversarial Networks (GANs) and Variational Auto-Encoders (VAEs) have been widely used for multiple data generation tasks and have obtained exceptional performances in previous studies [25, 26]. In both cases, the models learn a latent space to draw samples which cannot be discerned from real data. VAEs learn a latent space by solving the task of accurately reconstructing the original data using an encoder and a decoder [27], and GANs are unsupervised generative models based on a game theoretic scenario where two different networks compete against each other [28]. The synthetic data generated by these models can expand the diversity of samples that a model is trained on, potentially increasing the predictive performance and also improving the model generalization capabilities. It is important to emphasize that this comes at almost zero cost once the model is trained, contrary to generating new data. In addition, synthetic data has the advantage that, when it can serve as a faithful representation of real patient data, it can be easily shared without any regulatory hurdles for protected health information.

Several studies have focused on the generation of single-modality synthetic data for both RNA gene expression and WSI data. For example, the generation of gene expression data has been mainly in the context of data imputation and has been researched by leveraging the latent space of VAEs. Qiu et al. showed that *beta*VAEs, a special case of VAEs, can impute RNA-Seq data [29]. Similarly, Way et al. proposed a VAE trained on pancancer TCGA data, that is able to encode tissue characteristics in the latent space and also leverages biological signals [30]. Recently, Vinas et al. presented an adversarial methodology for the generation of synthetic gene expression profiles that closely resemble real profiles and capture biological information [31]. The generation of high-quality WSI tiles has also been researched in recent years given the success of GANs in generating natural images [32, 33]. For example, Quiros et al. showed that GANs are able to capture morphological characteristics of cancer tissues, placing similar tissue tiles closer in the latent space, while generating high-quality tiles [34, 35].

However, current research focuses on generating or imputing single modalities without leveraging the information provided from other data types. For non-medical applications, multi-modal data generation has made enormous progress thanks to the availability of large multi-modal data e.g. paired text and image data. Unsupervised learning methods such as GANs, transformers [36] and diffusion models [37] have been developed to leverage the relationship between these two modalities, enabling the generation of images based on their textual description [38–40], or generating textual descriptions of given images [41].

While multi-modal generation has proven successful for natural images in non-medical applications [38, 39], the relation between WSI and gene expression needs yet to be explored for multi-modal synthetic data generation. For this use case, we were inspired by the observation that the relation between textual descriptions and their corresponding images is similar to the relation between WSI images and genomic data, since they are describing the same phenomenon from two different perspectives.

Specifically, we explore the generation of WSI tiles using gene expression profiles of healthy lung and brain cortex tissue (Figure 1). First, we train a VAE that reduces the dimensionality of the RNA-Seq data. Then, using the latent representation of the gene expression as input, we present a GAN-based architecture that generates image tiles for healthy lung and brain cortex tissue. Based on evaluations of blinded pathologists, we show that the quality of generated WSI images can be significantly improved when the GAN is infused with gene expression data.

**Fig. 1.**
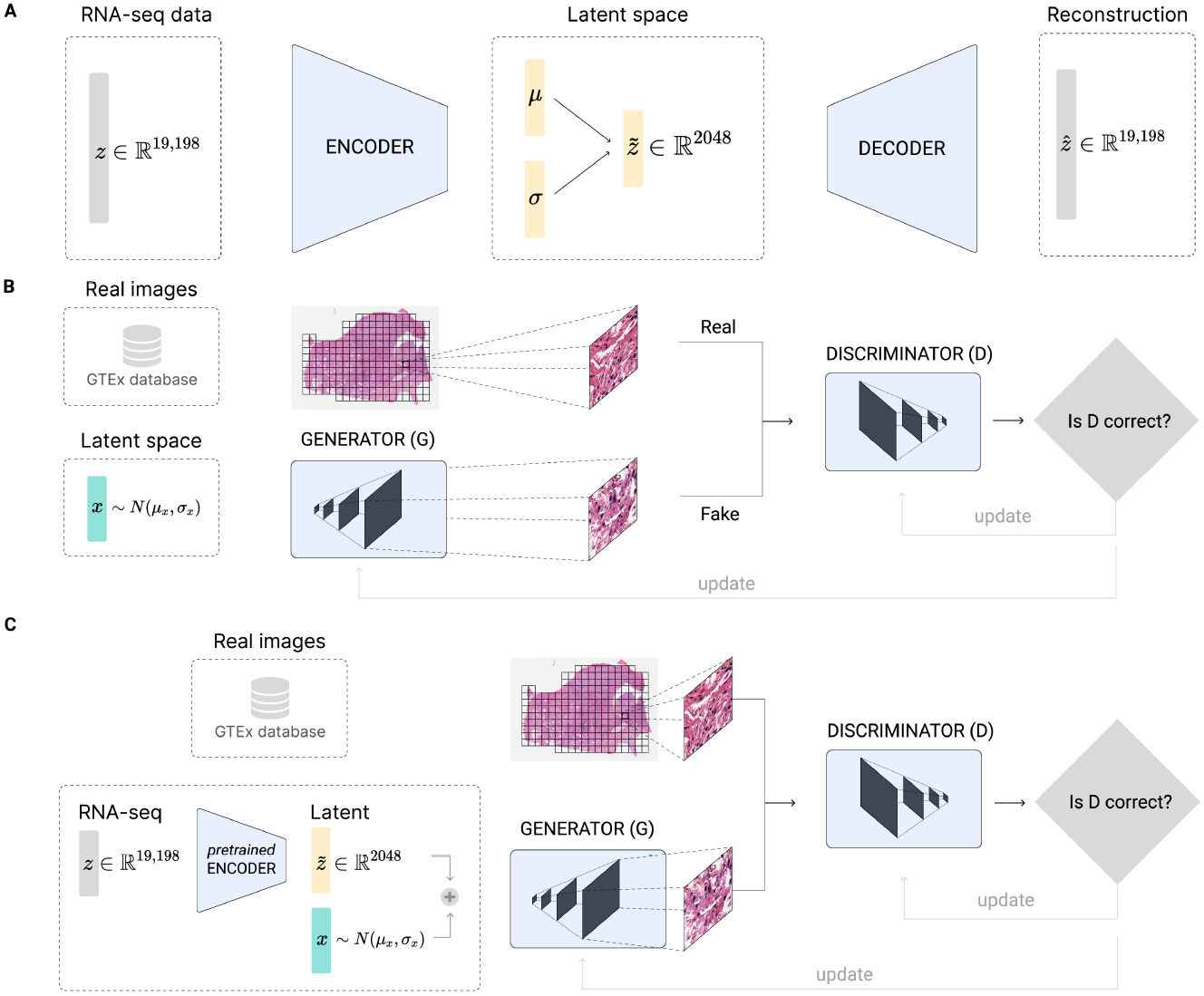
Model architecture for gene expression, WSI and combined data using VAE and GANs. **Panel A:** *β*VAE architecture for the generation of synthetic gene expression data. The model uses as input the expression of 19,198 genes. Both the encoder and the decoder are formed by two linear layers of 6, 000 and 4, 096 respectively. The latent *μ* and *σ* vectors have a feature size of 2, 048. **Panel B**: GAN architecture for generating tiles by sampling from a random normal distribution. The architecture chosen was a Deep Convolutional GAN (DCGAN) [42], using as input a feature vector of size 2, 048. The final size of the tiles generated is 256 × 256, the same as the size of the real tiles. **Panel C:** RNA-GAN architecture where the latent representation of the gene expression is used for generating tiles. The gene expression profile of the patient is used in the *β*VAE architecture to obtain the latent representation. Then a feature vector is sampled from a squeezed random normal distribution (values ranging between [-0.3, 0.3]) and added to the latent representation. A DCGAN is trained to use this vector as input and generate a 256 × 256. The discriminator receives synthetic and real samples of that size.

## Results

### A *β*-VAE model can build a representative latent space that discriminates between healthy tissues

As a first step, we aimed to create an accurate, distinguishable latent representation of healthy multi-tissue gene expression using a *β*-VAE architecture (Figure 1 A). The goal was to reduce the dimensionality of the gene expression profile while maintaining the differences among the tissues. To do so, we use the traditional approach for training a *β*-VAE, i.e. reconstructing the input from the latent space (see Methods). The *β*-VAE model was able to accurately reconstruct the gene expression by forwarding the latent representation through the decoder and obtaining a mean absolute error percentage of 39% (RMSE of 0.631) on the test set for multiple tissues.

To verify that the latent representation learnt by the *β*VAE accurately maps to the different tissues, the UMAP algorithm [43] was used to visualize the real gene expression data as well as reconstructions of latent representations on the test set. For lung and brain samples, two separated clusters can be distinguished, showing how the model is characterizing the two tissues in the latent space (Figure 2 A, ‘Real’ versus ‘Reconstruction’).

**Fig. 2.**
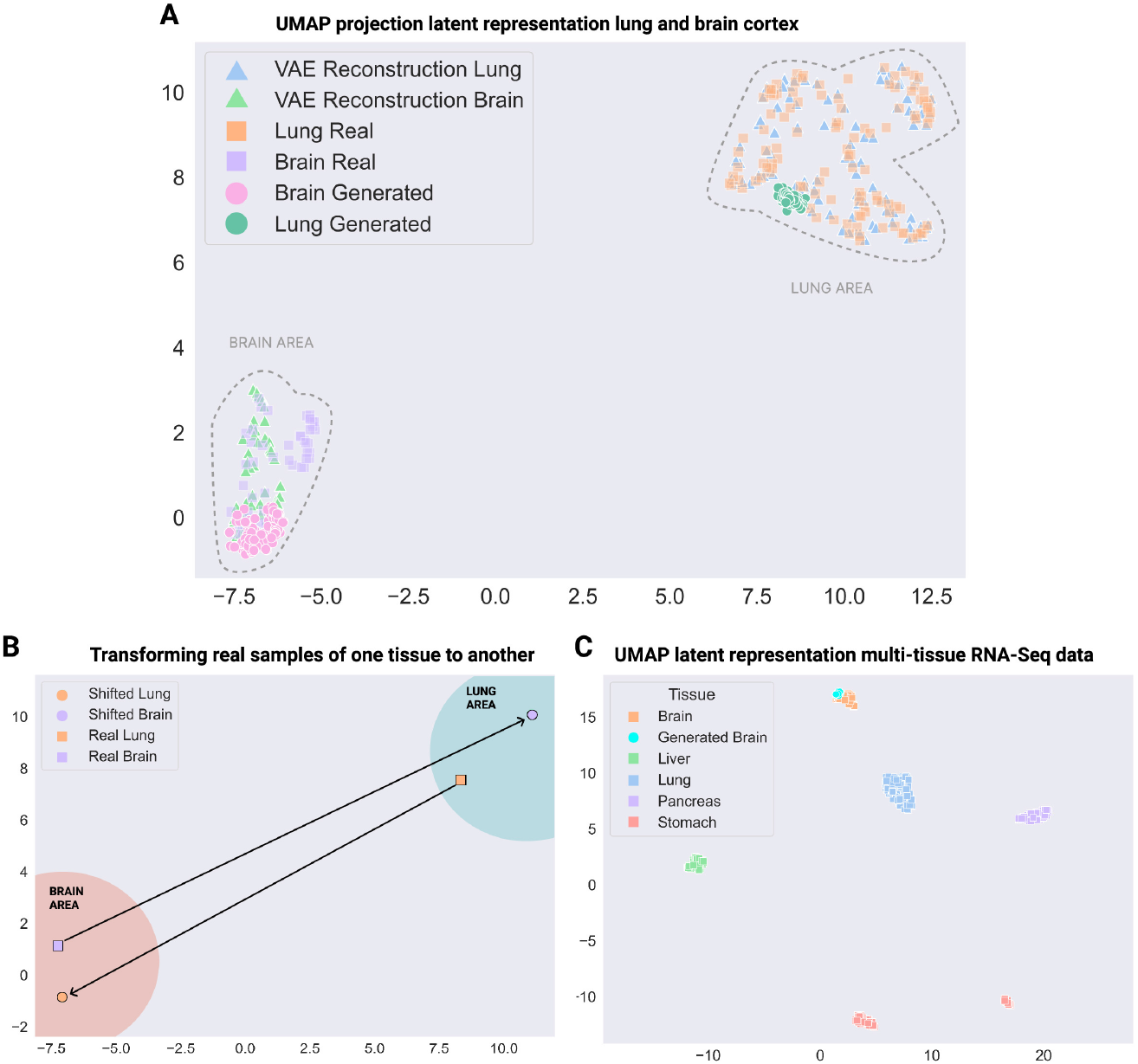
UMAP visualization of *β*-VAE embedding of multi-tissue expression profiles. **Panel A:** UMAP visualization of the real and reconstructed gene expression profiles of lung and brain cortex healthy tissue. Generated gene expression profiles, by sampling from the latent space and interpolating to the respective tissue, are also plotted. **Panel B:** Shifting real gene expression profiles between the two tissues. The latent representation of all the available samples is obtained, and the difference vectors between the cluster centroids are computed. **Panel C:** UMAP visualization of real gene expression profiles of multiple tissues and generated one from brain cortex tissue.

To further validate the learned latent space, we tested what happens when interpolating data in it. By interpolating in the latent space, we should be able to ‘transform’ a randomly drawn sample to a gene expression profile that looks like it originated from one of the tissues (i.e. synthetic gene expression generation). To do so, we need to calculate the cluster centroid vector over the real data latent representations of the desired tissue and add this centroid vector to randomly drawn samples from the *β*VAE latent distribution. This procedure allows us to generate synthetic gene expression data that look like real brain or lung gene expression data. When projecting these synthetic samples in the UMAP space, they indeed fall in the same clusters as the original data (Figure 2 A, ‘Generated’ versus ‘Real’).

We can also perform other operations in the latent space. For example, we should be able to ‘shift’ the gene expression from one tissue into what it would look like if it originated from another tissue. In this case, we need to add the difference vectors between the cluster centroids of the respective tissues to the latent representation of a given sample gene expression. For example, we can shift a real brain gene expression profile to a lung gene expression profile and vice versa. Visualizing these new samples in the UMAP space verifies that these operations can indeed be successfully performed (Figure 2 B). Next, the representation capabilities of the *β*VAE can also be extended to multiple tissues, showing a diverse representation with well-differentiated clusters, and maintaining the generative capabilities across the multiple tissues (Figure 2 C).

### GANs generate quality synthetic WSI tiles preserving real data distribution differences

Next, we developed a traditional GAN model to generate synthetic WSI tiles for brain cortex and lung tissue. The model was able to generate good quality images, preserving the morphological structures, and showing little artifacts (Figure 3 A, Supplementary Figure 1 A). In some tiles, checkerboard artifacts are noticeable, which is a known problem in GANs [44].

**Fig. 3.**
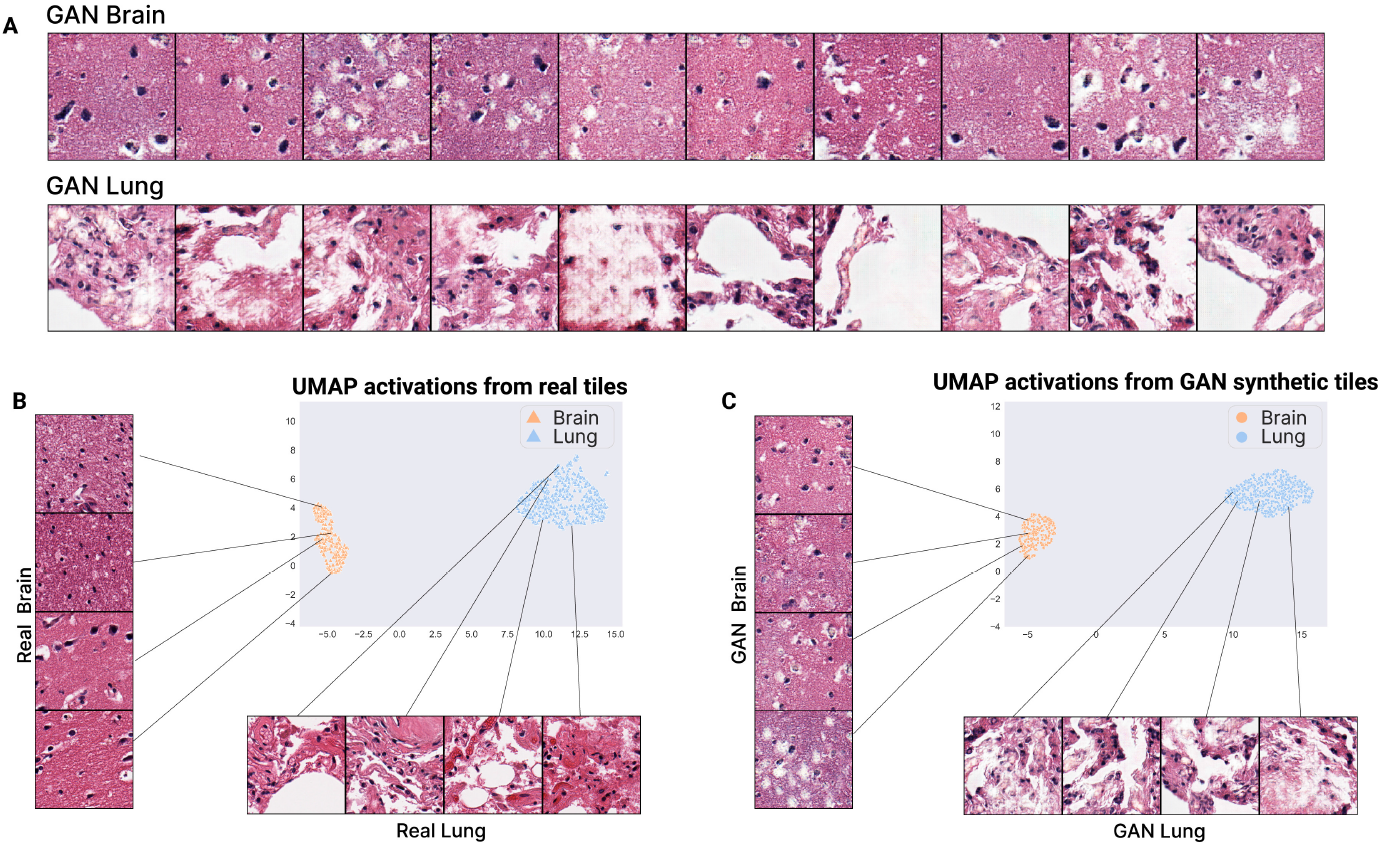
A GAN generates realistic lung and brain cortex tiles maintaining the distribution of the real tiles. **Panel A:** Tiles generated by the GAN model for brain tissue on the top and for lung tissue on the bottom. **Panel B:** UMAP representation of the real patients in the lung and brain cortex dataset. **Panel C:** UMAP representation of generated tiles using the GAN model. 600 tiles are generated per patient, and then used to compute the feature vectors and the UMAP visualization.

Despite the artifacts, the main cell types can be observed in the tiles, such as epithelial, connective, and muscle tissue. In addition, there is a clear distinction between the tiles generated for the brain cortex and the lung, preserving the characteristics of the corresponding real tiles. Specifically, the brain cortex tissue is grouped in a set of layers that form a homogeneous and continuous layer (the outer plexiform layer, outer granular layer, outer pyramidal cell, inner granular layer, inner pyramidal layer, and polymorphous layer) [45]. These characteristics can be observed in the synthetic brain tiles, i.e. they appear more homogeneous and contain less white spaces in comparison to the synthetic lung tissue tiles. The synthetic lung tissue tiles also present the characteristics of real tiles, showing the terminal bronchioles, respiratory bronchioles, alveolar ducts, and alveolar sacs in some cases.

To test if the generated tiles have the same distribution as the real ones, the feature vector outputted from one of the last convolutional layer of an Inception V3 network pretrained on Imagenet was obtained for the 600 generated tiles. Then, these feature vectors were projected and visualized using the UMAP algorithm, showing a similar distribution between the tissues for both real (Figure 3 B) and synthetic samples (Figure 3 C).

### Using latent gene expression profiles as input on GANs improves synthetic H&E tiles quality and reduces training time

Next, we used latent gene expression profiles as input instead of a random normal distribution for a GAN model generating WSI tiles. The gene expression was first forwarded through the pretrained *β*VAE to reduce the dimensionality and to encode it in the latent space. Then, that representation plus noise sampled from a narrowed random normal distribution (values between [-0.3, 0.3]) was used as input to the generator, that outputs the synthetic tile (Figure 1 C). This model generates synthetic tiles with fewer artifacts and better quality of the morphological structures (Figure 4 A, Supplementary Figure 1 B). To demonstrate that the gene expression latent representation provides actual information to generate the tiles and that the model does not mainly focus on the random signal (as with conventional GANs), we also created a GAN that samples only from a scaled random normal distribution (values between [-0.3, 0.3]). This model was not able to produce quality samples of any tissue (Figure 4 C). Hence, the RNA-Seq data in the RNA-GAN is the main signal that guides its sampling process.

**Fig. 4.**
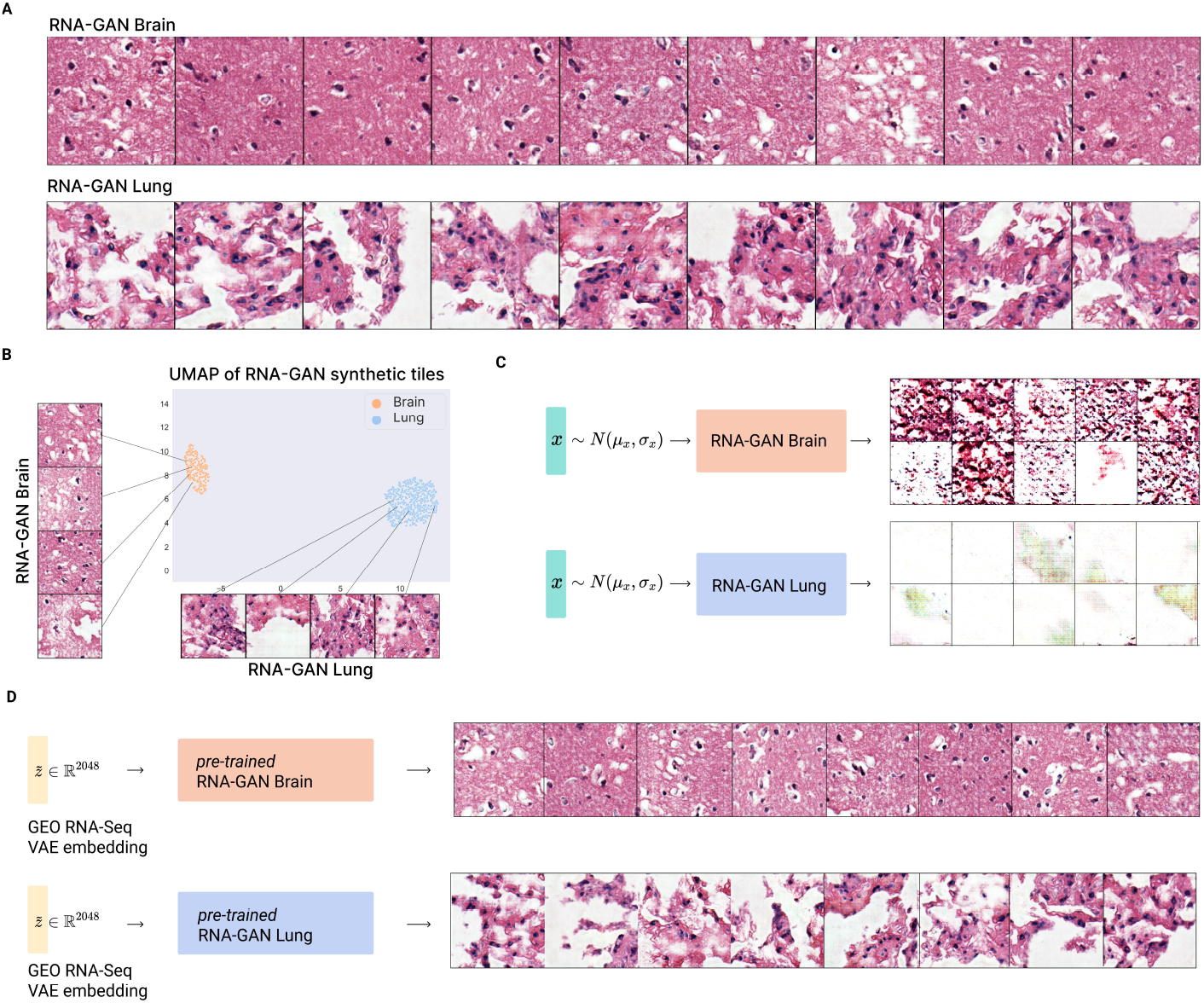
A gene expression infused GAN improves-tile quality. **Panel A:** tiles generated using the RNA-GAN model for lung and brain cortex healthy tissue. **Panel B:** UMAP visualization of the patients by generating tiles using their gene expression. The model preserves the distribution differences between the two tissues. **Panel C:** generated tiles of model trained using only random gaussian data on a small range ([-0.3, 0.3]) does not generate high quality tiles, showing that the gene expression distribution is essential for synthetic tile generation. **Panel D:** brain cortex and lung tissue tiles generated using an external data set (GSE120795), showing the generalization capabilities of the model.

We also obtained the feature vector from one of the last convolutional layers of the Inception V3 architecture pretrained on Imagenet, to observe if the distribution of the synthetic tiles was similar to that one from real patients. The differences between the tissues was preserved as well the tissue inner-cluster distribution (Figure 4 B).

To test the generalization capabilities of the trained models, we also used as input an external brain cortex and lung tissue RNA-Seq data (GEO series 120795). The model was able to successfully generate tissue samples with characteristics similar to those obtained with the training data (Figure 4 D). We then tested whether a model trained on real data can distinguish the synthetic generated tiles from this GEO cohort. This model reached an accuracy of 80.5%, a F1-Score of 79.7%, and an AUC of 0.805, showing that a model trained on real tiles can accurately classify the synthetic tiles. Finally, we observed that the RNA expression infused GAN model needed fewer training epochs in comparison to the regular GAN model (Figure 5).

**Fig. 5.**
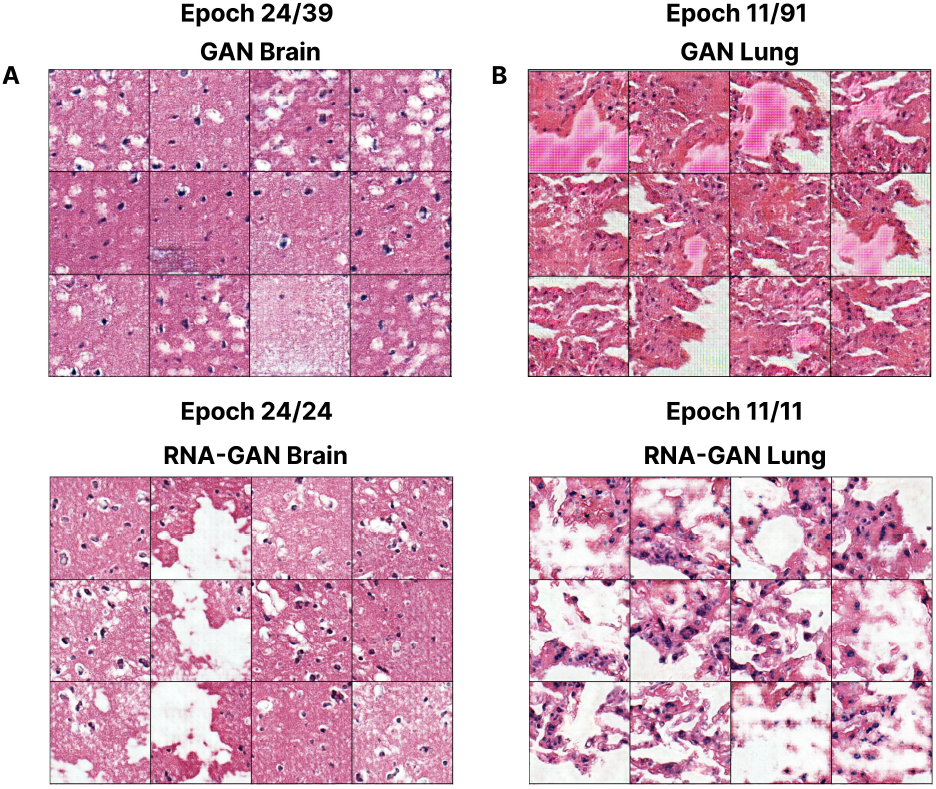
A gene expression profile-infused GAN converges faster: brain cortex and lung tissue tiles generated at the same epoch during training for the model with and without gene expression profiles. The visualized epoch is the last epoch of training for the models using RNA-Seq data. **Panel A:** brain cortex generation at training epoch 24 for GAN and RNA-GAN models, with similar performance and quality between the generated tiles, however, less diversity is obtained when not using gene expression profiles. **Panel B:** lung tissue generation at training epoch 11 for both the GAN and RNA-GAN models. A comparison of both models show noticeable differences in the quality of the generated tiles. The model using gene expression profiles outputs better morphological features, less artifacts, and has a higher overall quality.

### Expert evaluation of synthetic tiles

Next, we asked a panel board-certified anatomic pathologists with different sub-specialty expertise (MO, HV, RBW, CSK, MVR, JS) to rate the quality of brain cortex and lung cortex tiles (N=18), on a scale from 1 (worst) to 5 (best). These pathologists were not informed about the presence of synthetic data in the examples.

The pathologists’ evaluation of the morphological structures resulted in a mean score of 3.55 ± 0.95 for real brain, 2.88 ± 0.62 for GAN brain, and 2.94 ± 0.64 for RNA-GAN brain. For the lung tissue, the mean score for the real samples was 2.26 ± 1.14, 1 ± 0.55 for the GAN lung, and 1.73 ± 0.79 for the RNA-GAN lung. Hence, the pathologists rated the real samples as best quality, with second-best ratings for the samples from the RNA-GAN and the worst ratings for those from the conventional GAN.

The preference in the pathologists’ evaluation for the RNA-GAN lung over GAN synthetic tiles is statistically significant (*p – value* = 0.025), while there is no statistically significant difference in ratings between the real and RNA-GAN lung tiles scores (*p – value* = 0.052). Further, a bigger mean evaluation score difference was found between real and GAN tiles than between real and RNA-GAN tiles, again confirming that the quality of RNA-GAN synthetic tiles is closer to the quality of real tiles (Figure 6 A). In addition, the mean difference in the evaluation between GAN and RNA-GAN tiles was bigger than zero, showing the preference of pathologists for the RNA-GAN tiles over GAN-tiles (Figure 6 B).

**Fig. 6.**
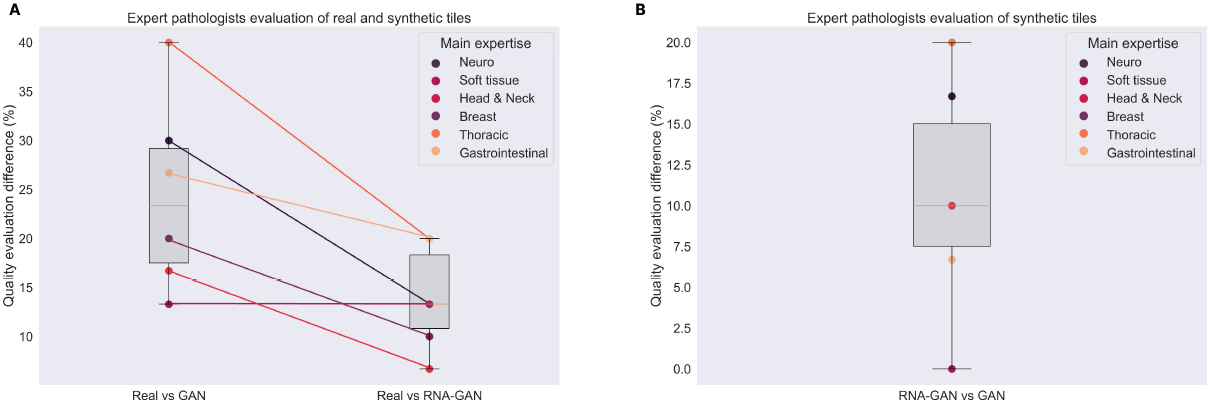
Expert evaluation of synthetic slides. **Panel A:** difference in morphological structure quality of synthetic (generated by GAN and RNA-GAN) and real tissues based on the pathologists’ evaluation. The difference between real tiles and generated tiles was bigger for GAN than for RNA-GAN. **Panel B:** Difference in morphological structure quality between the synthetic generated tiles by the GAN and RNA-GAN based on the pathologists’ evaluation. Pathologists evaluated the tiles generated using RNA-GAN better in comparison to only GAN.

Finally, pathologists detected the tissue of origin of the tiles with a 100% ± 0.0 accuracy for both real and RNA-GAN tiles, while this drops to 74.98% ± 20.40 accuracy for the GAN tiles.

## Discussion

Since biomedical data is becoming increasingly multi-modal, there is a growing interest in developing multi-modal predictive models for advancing the goals of precision medicine. However, obtaining multi-modal biological data is a slow and costly process. Multi-modal data is sometimes available for widely studied diseases, but this is not common and definitely not the case for pediatric and rare diseases. In addition, not all medical facilities have the required expertise or instruments to collect each data modality for a patient. Therefore, multimodal data imputation [29, 46, 47] and generation of multi-modal synthetic data is a promising approach to complete data sets by leveraging the potential of deep learning models [24].

Previously, imputed samples and synthetic data were generated in an isolated way, not using the information provided by other modalities. In contrast, in this work, we studied the generation of WSI tiles of lung and brain cortex tissues by leveraging the corresponding gene expression profiles. This idea was inspired by recent advances in multi-modal data generation for nonmedical images, including the generation or modification of images based on text prompts (e.g. DALL-E 2 [39], Imagen [48], Parti [49] and related models [38, 39, 41]). Here, we extrapolated this idea to biomedical data by treating RNA-sequencing data as prior text to contextualize image generation [50, 51].

As a first step in our work, we developed a *β*VAE model that reduces the dimensionality of gene expression data. We have shown that the *β*VAE model is able to capture the latent representation of multiple human tissues, and that it obtains a representative latent feature vector that accurately distinguishes between the tissues (Figure 2). To test the accuracy, practicality and capabilities of the learned latent space, we performed certain sanity checks. For example, we tested the ability to generate synthetic gene expression profiles and to interpolate samples between classes. Our analysis shows that the *β*VAE model indeed allows to obtain a compact, accurate representation of tissue gene expression profiles. We later use the encoder part of the *β*VAE to reduce the dimensionality of gene expression profiles which in turn will guide the generation of the WSI tiles. Note that this compact representation could also be used for downstream tasks including prognosis or treatment outcome prediction, but this analysis was out of the scope of this work.

Next, we trained a traditional GAN model on brain cortex and lung tissue data to generate WSI tiles. We explored the possibility of using the same architecture to generate both tissues, but the model collapsed, and only generated brain cortex tiles. We hypothesize this could be due to the homogeneity of brain tissue compared to lung tissue, making it easier for the model to generate brain cortex tiles and reduce the loss. This can also be observed in the number of necessary epochs to generate quality tiles for each tissue. While the model only needs 34 epochs for training convergence for brain cortex tissue, it requires 91 epochs for lung tissue. Clearly, it is more difficult to generate the lung tiles, probably because lung tissues are more heterogeneous compared to brain cortex. Importantly, the synthetic tiles preserved the distribution of real tiles, showing two well-differentiated clusters using a UMAP projection of the feature vectors (Figure 3 C). Pathologists where asked if they could observe any kind of artifact on the synthetic tissue (e.g. image aberrations). From the presented GAN tiles, in 70% of cases, pathologists detected certain artifacts (mean percentage across pathologists), while this was only the case for 17% of the real tiles.

Finally, we trained the gene expression profile infused GAN model, both on brain cortex and lung tissue data. The quality of the generated tiles improved significantly in comparison to tiles from a regular GAN, based on evaluation of expert pathologists. In addition, pathologists reported significantly fewer artifacts in RNA-GAN images (56% in comparison to 70% for GAN images). Another advantage of the RNA-GAN model over the GAN model is that it needed less training epochs for reaching high-quality results (using the same amount of training tiles). While the GAN lung model was trained during 94 epochs for obtaining tiles without major artifacts and good tissue structure, the RNA-GAN lung model only needed 11 epochs. This reduces the number of training epochs by 88%.

Once we have a trained generative model, generating or imputing missing values with synthetic data comes at very low cost. Adding these low-cost synthetic samples to real data can help to train more complex deep learning models which are data-hungry. While the synthetic generation of WSI and imputation of RNA-Seq data has been studied in the literature [29, 30, 34], to our knowledge generation of synthetic WSI tiles using gene expression profiles has not yet been explored. Here, we showed that infusing GANs with gene expression data for WSI tile generation does not only increase the quality of the resulting tiles, but additionally requires fewer compute time compared to the regular, single-modality approach.

We further showed how our model was able to generalize to gene expression profiles outside of the training dataset, still generating realistic synthetic tiles. While a vast amount of gene expression datasets are publicly available, most of these datasets do not provide matching WSIs. This limits their use in multimodal classification models [13, 15, 20, 52]. With our model, WSI tiles can be imputed from the gene expression, which opens new possibilities in publicly available datasets.

In summary, our work shows the promise of multi-modal synthetic biological data generation to obtain better quality multi-modal data for training complex, data-hungry deep learning models. This is especially useful for multimodal problems [13, 15] and datasets with missing modalities, but also for problems with small datasets restricted by expensive data collection or rare diseases. In future work, we intend to extend this approach to more heterogeneous tissues and complex diseases. We will also explore other GAN architectures [32, 33, 53] and diffusion models [37, 54].

## Methods

### Data

Data was obtained from The Genotype-Tissue Expression (GTEx) project [55]. The GTEx project aims to build a public resource of healthy tissue-spefic characteristics, providing gene expression and WSI among other data types. The data was collected from 54 non-disease tissue and across almost 1, 000 individuals. We collected the RNA-Seq and WSIs from brain cortex, lung, pancreas, stomach and liver tissues. There were a total of 246 samples of brain cortex tissue, 562 samples of lung tissue, 328 samples of pancreas tissue, 356 of stomach tissue, and 226 samples of liver tissue. To validate the generalization capabilities in generating tiles from the gene expression of other cohorts, the GEO series 120795 was used [56].

### RNA-Seq data preprocessing

Gene expression data from the GTEx project contains a total of 56, 201 genes. This number would require huge computational capabilities, and it difficulties the training of the machine learning models. Therefore, we reduced the feature dimension and obtained the expression of 19, 198 protein coding genes for further experiments. The data was log transformed, and the z-score normalization was applied to the gene expression using the training set data, in order to not include the validation or the test set information on the normalization process.

In the generalization experiments, the gene expression from lung and brain cortex tissue of the GEO series 120795 was used. However, not all the previously selected protein coding genes where among those sequenced in this dataset. Therefore, for those missing in this external cohort, we initialized them as zero for the generation of the tiles. Data was normalized using the mean and standard deviation from the training set of the GTEx data and log transformed.

### WSI data preprocessing

GTEx WSIs were acquired in SVS format and downsampled to 20 × magnification (0.5*μm* px^−1^). The size of WSIs is usually over 10*k* × 10*k* pixels, and therefore, cannot be directly used for training machine learning models to generate the data. Instead, tiles of a certain dimension are taken from the tissue, and they are used to train the models, which is consistent with related work in state-of-the-art WSI processing [5, 57, 58]. In our work we took non-overlapping tiles of 256 × 256 pixels. Firstly, a mask of the tissue in the higher resolution of the SVS file was obtained using the Otsu threshold method [59]. Tiles containing more then 60% of the background and with low-contrast were discarded. A maximum of 4, 000 tiles were taken from each slide. For the pre-processing of the images we relied on the python package openslide [60], that allows to efficiently work with WSI images. The tiles were saved in a LMDB database using as index the number of the tile. This approach enables to reduce the number of generated files, and structure the tiles in an organized way for a faster reading while training.

### *β*VAE architecture for synthetic gene expression generation and experiments

We chose the *β*VAE model for the generation of synthetic gene expression data [61]. The *β*VAE model is an extension of the VAE where a *β* parameter is introduced in the loss function. The original auto-encoder is formed by two networks, the encoder and the decoder. The encoder encodes the input into a lower dimensionality representation, and then it is used to reconstruct the input using the decoder, by learning the function *h_θ_*(*x*) ≈ *x* being *θ* the parameters of the neural network. To learn this function, we want to minimize the reconstruction error between the input and the output. The most common loss function is the root mean squared error (RMSE). However, for the VAE we want to learn a probability distribution of the latent space, which allows to later sample from it to generate new samples. The assumption of the VAE is that the distribution of the data x, *P*(*x*) is related to the distribution of the latent variable *z, P*(*z*). The loss function of the VAE, which is the negative log-likelihood with a regularizer is as follows:

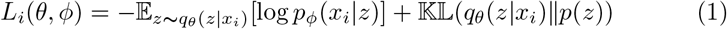

where the first term is the reconstruction loss and the second term is the Kullback-Leibler (KL) divergence between the encoder’s distribution *q_θ_*(*z|x*) and *p*(*z*) which is defined as the standard normal distribution *p*(*z*) = *N*(0,1).

For the *β*VAE we introduce the parameter *β*, which controls the effect of the KL divergence for the total loss:

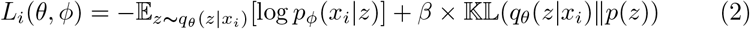

If *β* = 1, we have the standard loss of the VAE. If *β* = 0, we would only focus on the reconstruction loss, approximating the model to a normal autoencoder. For the rest of the values, we are regularizing the effect of the KL divergence on the training of the model, making the latent space smoother and more disentangled [61].

For the final architecture, we empirically determined to use two hidden layers of 6, 000 and 4,096 neurons each for both the encoder and the decoder, and a size of 2,048 for the latent dimension. Given that we were going to use the latent representation for the generation of the tiles, we followed the same dimensionality as the output of the convolutional layers of state-of-the-art convolutional neural networks [62]. We used batch norm between the layers and the LeakyReLU as the activation function. A *β* = 0.005 was used in the loss function. We used the Adam optimizer for the training with learning rate equal to 5 × 10^−5^, along with a warm-up and a cosine learning rate scheduler. We trained the model for 250 epochs with early stopping based on the validation set loss, and a batch size of 128. A schema of the architecture is presented in Figure 1 A.

We divided the dataset in 60-20-20 % training, validation and test stratified splits. We trained two different models, one for brain cortex and lung tissue data, and the other with all the tissues described in previous subsections (lung, brain cortex, stomach, pancreas, and liver).

### GAN architecture for synthetic WSI tiles generation and experiments

GANs have been succesfully used for generating high-fidelity images. In this work we use the Deep Convolutional GAN architecture presented by Radford et al. [42]. On early experiments we used the minmax loss function described on the original Godfellow et al. [28] work. However, this loss function led to a lack of diversity in the generation of the samples, and a diminished quality. Therefore, we decided to use the Wasserstein loss introduced by Arjovsky et al. [63], also adding the gradient penalty proposed by Gulrajani et al. [64]. In this case the discriminator (or critic, as called in the paper) does not classify between real and synthetic samples, but for each sample it outputs a number. The discriminator training just tries to make the output bigger for real samples and smaller for synthetic samples. This simplifies the loss function of both networks, where the discriminator tries to maximize the difference between its output on real instances and its output on synthetic instances as follows:

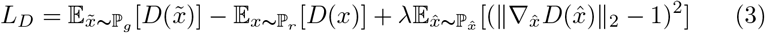

and the generator tries to maximize the discriminator’s output for its synthetic instances as follows:

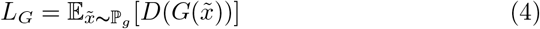

We trained two DCGANs, one per tissue, by sampling from a normal random distribution (scheme depicted in Figure 1 B). We samples different number of tiles per image for the training of the network, finally selecting 600 tiles per image because the quality of the image and the artifacts were highly improved by augmenting the number of tiles. We used the Adam optimizer for both the generator and the discriminator, with a learning rate equal to 1 × 10^−3^ for the generator, a learning rate equal to 4 × 10^−3^ for the discriminator and betas values (0.5,0.999) in both cases. Data augmentations such as color jitter and random vertical and horizontal flips were used during training. The brain tissue GAN was trained during 39 epochs while the lung tissue GAN was trained during 91 epochs. For the training of the GANs, the Python package Torchgan was used [65].

### RNA-GAN architecture for synthetic WSI tiles generation and experiments

After the successful generation of the tiles using a traditional GAN approach, we explored the generation of synthetic tiles by using the gene expression profile of the patient. We combined the pretrained *β*VAE with the DCGAN architecture, using the encoding in the latent space as the input for training the generator. To generate different tiles from the same gene expression profile, we sample a noise vector from a narrowed random normal distribution (values ranging between [-0.3, 0.3]) and add it to the latent encoding. Therefore, the input to the generator would be:

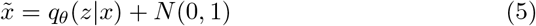

We trained two DCGANs, one per tissue, and the pipeline is depicted in Figure 1 C). We finally selected 600 tiles per image to train the generator. We used the Adam optimizer for both the generator and the discriminator, with a learning rate equal to 1 × 10^−3^ for the generator, a learning rate equal to 4× 10^−3^ for the discriminator and betas values (0.5, 0.999) in both cases. Data augmentations such as color jitter and random vertical and horizontal flips were used during training. The brain tissue GAN was trained during 24 epochs while the lung tissue GAN was trained during 11 epochs. For the training of the GANs, the Python package Torchgan was used [65].

To validate the generalization capabilities of the trained model, the GEO series 120795 was used. It contains gene expression profiles from healthy tissues, were we took the expression of lung and brain cortex tissues. For obtaining machine learning performance metrics, one hundred images were generated per tissue. Then, a Resnet-18 was trained from scratch using 10 epochs and early stopping based on a 20% of data as validation set. A learning rate value of 3*e*^−5^ and AdamW optimizer were used. Finally, the model was tested on the GEO synthetically generated data, and accuracy, F1-Score and AUC was computed.

We release an online quiz where users can try to distinguish between real and synthetic samples, obtaining a score on how well they performed. The quiz is available in the following URL: https://rna-gan.stanford.edu/. The code and checkpoints for the proposed models is available in this Github repository: https://github.com/gevaertlab/RNA-GAN.

### Expert pathologist evaluation form

To evaluate the quality of the synthetic tiles, we presented a form to expert pathologists (Supplementary Material Figure 2). The pathologists were not informed that some presented tiles were synthetic, to omit any kind of biases on the evaluation. Instead, we informed the pathologists that these tiles were going to be used to create machine learning classifiers, and we wanted to evaluate their quality for this task. Three questions were asked to the experts:

1. Is the tile from brain cortex or lung tissue?
2. Quality of the morphological structures: Being 1 very bad and 5 very good, how would you rate the morphological features present in the tile for an assessment of the tissue?
3. Do you find artifacts in the image? (e.g. image aberrations) (Yes/No)

---

**Supplementary Figure 1.**
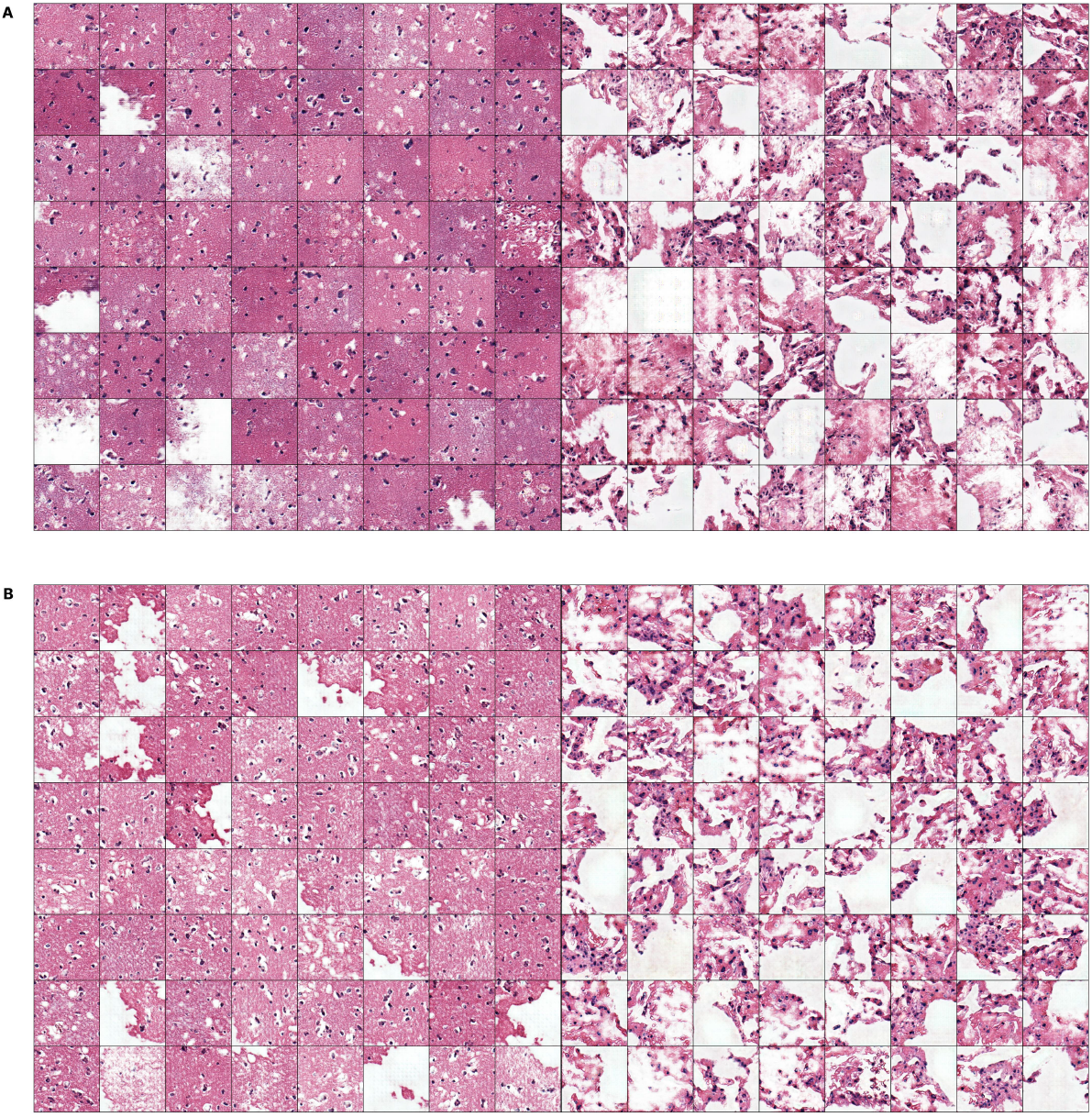
**Panel A:** More examples of brain cortex and lung tissue generation using the GAN architecture. **Panel B:** More examples of brain cortex and lung tissue generation using the RNA-GAN architecture.

**Supplementary Figure 2.**
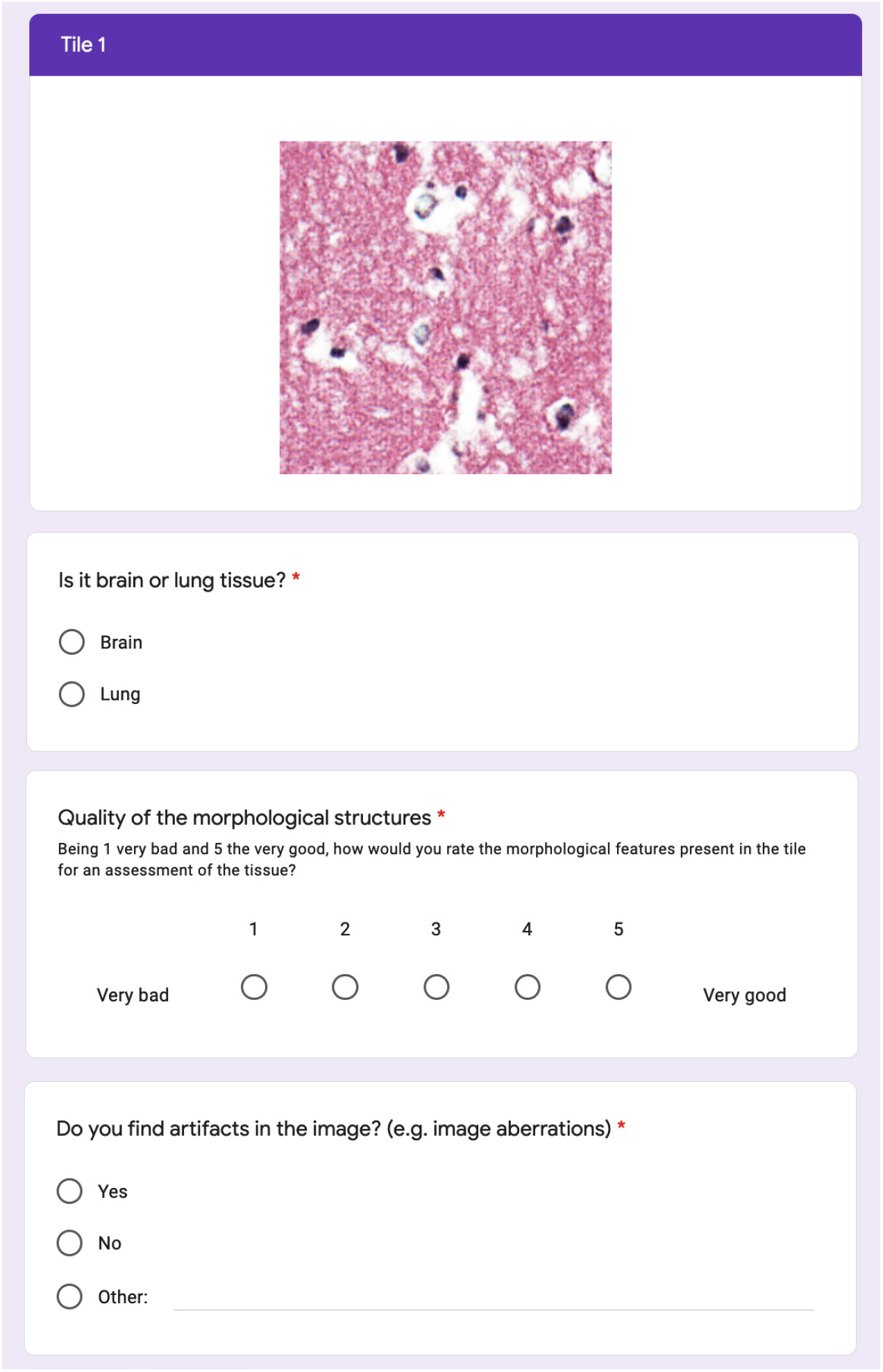
Example of one of the samples of the form presented to expert pathologists. Three question were asked, regarding the tissue of origin, the quality of the morphological structure and the presence of image artifacts.

## Acknowledgments

We would like to thank Matt Van den Rijn for his collaboration evaluating the synthetic images. FCP and LJ were supported by the Spanish Ministry of Sciences, Innovation and Universities under Projects RTI-2018-101674-B-I00 and PID2021-128317OB-I00, and the project from Junta de Andalucia P20-00163. FCP was also supported by a Predoctoral scholarship from the Fulbright Spanish Commission. Research reported here was further supported by the National Cancer Institute (NCI) under award: R01 CA260271. The content is solely the responsibility of the authors and does not necessarily represent the official views of the National Institutes of Health. MP was supported by a fellowship from the Belgian American Educational Foundation.

## Author contributions

Conceptualization, F.C.P. and O.G.; Methodology, F.C.P, M.P., and O.G.; Investigation, F.C.P., M.P., M.O., H.V., R.W., C.K., L.J.H., J.S., and O.G.; Writing – Original Draft, F.C.P, M.P., and O.G.; Writing-Review & Editing, M.O., H.V., R.W., C.K., L.J.H.,and J.S.; Funding Acquisition, O.G. and L.J.H.; Resources, O.G.; Supervision, O.G.

## Declarations

The authors declare no competing interests.

